# Shox2 is necessary for normal thalamic spindle function

**DOI:** 10.1101/2024.09.10.612202

**Authors:** Isabella Febbo, Matthieu Maroteaux, Diankun Yu, Valerie Warkins, LA Martinez, AE Anderson, MJ Galazo, LA Schrader

## Abstract

The cellular identity of thalamocortical neurons (TCNs), namely their firing properties, dictates brain-wide activity patterns, such as sleep spindles. Transcription factors are critical to the determination of cellular identity. Previously, we discovered that a subset of TCNs express the transcription factor, Shox2, and, in a global *Shox2* KO, established that TCNs within the anterior nucleus of the thalamus rely on the expression of *Shox2* to regulate key ion channels that are necessary to maintain their firing properties. From this, we hypothesized that *Shox2* expression, through the regulation of firing properties of TCNs, is critical for the thalamocortical circuit to generate spindle oscillations. We exploited the somatosensory thalamocortical circuit to investigate this by creating a primary somatosensory thalamus (VB) *Shox2* knockdown mouse model. We delivered Cre into the VB of P21 Shox2^fl/fl^ mice using viral infection and compared *in vitro*, patch-clamp recordings from *Shox2*^+^ and *Shox2* knockdown TCNs, finding that *Shox2* expression is indeed critical to maintain burst and tonic firing properties of VB TCNs. Since Shox2 is important developmentally and firing from TCNs to cortex during development structures the circuit, we performed ultrasound-guided P3 injections at P3 to generate an early-stage, Shox2 VB knockdown, but found no changes in the layer four, barrel map (VB cortical target). Despite this, Shox2 knockdown mice exhibit reduced sleep-spindle EEG density. Further, key behaviors associated with spindles and proper VB thalamic function—memory consolidation and somatosensory perception—are significantly impaired. These results indicate that the impact on spindle function is likely due to cell autonomous changes to TCNs rather than circuit changes, confirming our hypothesis that *Shox2* is necessary for normal thalamic spindle function and implicating a potential role for *Shox2* in autism and schizophrenia pathologies.

## Introduction

The circuitry and firing activity of neurons, mediated by an interplay between network connectivity and intrinsic cellular properties, underlie neural computations. Properties of cells and their connections contribute to the emergence of rhythms in the brain that are critical to the processing of cognitive and sensory information. These rhythms, or oscillations, occur at different frequencies and contribute to unique neural functions. For example, beta oscillations from 12 to 20 Hz in the parietal cortex integrate multimodal information to generate a representation of recent sensory inputs (Gelastopoulos, Whittington et al. 2019). Another example is the sleep spindle, occurring int the thalamocortical loop at 7 to 14 Hz, these oscillations are important in memory consolidation during sleep (Latchoumane, Ngo et al. 2017). Interestingly, reduced density of sleep spindles is a recently determined biomarker for autism and schizophrenia (Manoach, Pan et al. 2016, Farmer, Chilakamarri et al. 2018, Copping and Silverman 2021).

Sleep spindles originate in the thalamus, as they are generated by firing between interconnected thalamocortical and thalamic reticular neurons (Steriade, Domich et al. 1987, Astori, Wimmer et al. 2011, Halassa, Siegle et al. 2011). Thalamocortical neurons (TCNs) then relay these thalamus-generated spindles to the cortex, where they can be recorded on electroencephalograms (EEGs). Since sleep spindles are generated by the thalamus, they are a valuable indicator of proper thalamic activity and thalamocortical signal relay to the cortex. Generation of these spindles, along with all oscillatory thalamic behavior, relies upon precisely timed burst and tonic firing of TCNs (Kandel and Buzsaki 1997, Halassa, Siegle et al. 2011, Lee, Song et al. 2013). TCNs can fire in burst or tonic modes (Guido and Weyand 1995, Sherman 2001), and proper cortical receipt of this thalamic information relies critically upon the timing and synchrony of this firing (Bruno 2011), as demonstrated *in vitro* and *in vivo* (Sherman 2001, Swadlow and Gusev 2001, Swadlow 2002, Lesica and Stanley 2004, Lesica, Weng et al. 2006, Wang, Webber et al. 2010, Stanley, Jin et al. 2012, Whitmire, Waiblinger et al. 2016, Borden, Wright et al. 2022). This signal is then returned to TCNs via corticothalamic projections as a modulatory input, which elicits burst firing of TCNs and generates thalamocortical oscillations that underlie higher order thalamic functions, such as the sleep spindles which are important for memory consolidation (Gais, Molle et al. 2002, Latchoumane, Ngo et al. 2017). As such, the TCN membrane properties that regulate TCN firing are foundational to sensory perception and thalamocortical-related cognition.

TCNs are intrinsic oscillators that burst fire in response to membrane hyperpolarization and fire tonically when depolarized. These properties are generated by TCN membrane expression of specific ion channels, notably, T-type Ca^2+^ and HCN channels, and subtypes of potassium channels (Espinosa, Torres-Vega et al. 2008, Amarillo, Zagha et al. 2014). T-type Ca^2+^ and HCN channels mediate TCNs’ synchronous burst firing in response to hyperpolarization (Jahnsen and Llinas 1984, Luthi, Bal et al. 1998), while potassium channels are critical to timing of tonic firing in response to depolarization (Rudy and McBain 2001, Kasten, Rudy et al. 2007). Thus, understanding the mechanisms that control these firing properties is important to overall thalamic function.

Cell identity, including the distribution of channel types and consequent firing properties, are strongly determined by transcription factors. Despite the critical role of the thalamus in cognition, relatively little is known about the transcription factors that regulate maturation and maintenance of thalamic neuron membrane properties. Most investigation into the function of thalamic transcription factors examine embryonic and developmental determinants of thalamic nuclei organization (Nakagawa and O’Leary 2001, Nakagawa and O’Leary 2003, Yuge, Kataoka et al. 2011). In the adult mouse thalamus, combinatorial expression of *Tcf7l2, Lef1, Gbx2, Prox1, Pou4f1, Esrrg*, and *Six3* define molecular divisions that loosely correlate with established nuclei divisions (Nagalski, Puelles et al. 2016). Pax6 and Gli2 have been identified as transcription factors that are critical to the development of thalamocortical projections (Pratt, Vitalis et al. 2000, Callejas-Marin, Moreno-Bravo et al. 2022). TCF7L2, is expressed throughout the thalamus as well as the habenula, and is critical for thalamocortical projections and habenula organization (Bem, Brozko et al. 2019). These developmental studies indicate proper determination and regionalization of the thalamus; however, no studies have investigated factors that are necessary for generation or maintenance of TCN firing properties.

Shox2 is expressed in both human and mouse thalamus (Nagalski, Puelles et al. 2016), and it is involved in the regulation of rhythmic firing properties in the heart, spinal cord, and thalamus (Puskaric, Schmitteckert et al. 2010, Dougherty, Zagoraiou et al. 2013, Sun, Yu et al. 2015, Yu, Febbo et al. 2021). Our previous study investigated the role of Shox2 in neuron intrinsic and firing properties in a higher order nucleus associated with cognitive function, the anterior nucleus of thalamus. Here, we validate our previous findings that Shox2 expression is critical for the firing properties of TCNs in the ventrobasal (VB) nucleus of the thalamus using a VB targeted KO model. We analyzed the effect on TCN burst firing, tonic firing, membrane properties and cortical target organization. We then further determined if thalamocortical oscillations and thalamic-related behavior were disrupted in *Shox2* KO mouse models. We found that *Shox2* expression is critical to membrane and firing identity of TCNs, as well as the emergent thalamocortical oscillations. Spindle density is significantly reduced in *Shox2* VB KO, and memory consolidation and sensory perception are impaired. These experiments provide evidence that the transcriptional activity of Shox2 generates TCN firing characteristics that promote proper thalamic function, specifically: sensory perception and memory consolidation.

## Results

### Shox2+ TCNs possess typical connectivity with cortex and reticular thalamus

Our previous study demonstrated *Shox2* expression in TCNs (Yu et al. 2021)Click or tap here to enter text.. The present study investigates the cortical and reticular thalamus connectivity of *Shox2*-expressing (*Shox2+*) TCNs. The VB nucleus of the thalamus receives somatosensory information and consists of first order TCNs, and *Shox2* is highly expressed in these neurons. First order TCNs are a subset of TCNs that are located throughout the thalamus and are generally associated with receiving primary sensory inputs from the brainstem and sending information on to the cortex. The barrel cortex is the well-defined area of the somatosensory cortex that receives and processes sensory information from the whiskers on the mystacial pad of the snout. First order TCNs of VB project to the somatosensory barrel cortex and to the reticular nucleus of the thalamus (Jones 2002). In the barrel cortex, they project to all layers, but anatomically most densely to layers IV and VI (Hersch and White 1981, Agmon and Connors 1991, Oberlaender, Ramirez et al. 2012). Functionally, this is supported by studies where optogenetic stimulation of VB projections evokes responses from layer IV that are significantly larger than any other layer. VB TCNs project mainly to excitatory and PV+ inhibitory cells in cortical layer IV within the barrels (Chmielowska, Carvell et al. 1989, Wimmer, Bruno et al. 2010, Sermet, Truschow et al. 2019) due to the abundance of *Shox2* + neurons located in the largely first order VB nucleus, we hypothesized that *Shox2*+ TCNs also make these first order TCN connections.

VB TCNs also synapse on the reticular nucleus (Jones 2002), which is an inhibitory nucleus surrounding the thalamus and is comprised of GABAergic interneurons. These thalamo-reticular connections are critical for sleep spindle generation. Sleep spindles are abolished in TCNs that are disconnected from the reticular nucleus (Steriade, Deschenes et al. 1985). However, the reticular nucleus in isolation from the dorsal thalamus, continues to generate spindle rhythms (Steriade, Domich et al. 1987).

To confirm these synaptic connections and visualize *Shox2^+^* cell somas and their projections, we investigated *Shox2* expression in the *Shox2-*Cre mouse crossed to the Ai14 reporter mouse, which expresses red fluorescent protein (RFP) in Cre*+* neurons. A sagittal image from a Shox2-Ai14 offspring (**figure 1A**) reveals localization of strong *Shox2* expression in the thalamic neurons with projections to various layers of the cortex including cortical layers IV and VI, as well as projections to the reticular nucleus. To visualize synaptic connectivity in cortex layer IV (**top**), layer VI (**middle**), and reticular nucleus (**bottom**) (**figure 1C**), we utilized the reporter mouse to visualize projections of *Shox2*-expressing TCNs in red (**column 2**), stained for glutamatergic synapses, using a Shank2 antibody—in far red (**column 3)**, and PV+ interneurons—using a parvalbumin antibody—in green (**column 4)**. Merging these three wavelengths (**column 5**) results in visually identifiable synapses at high resolution, indicated by white arrows. The PV+ interneurons in the cortex and reticular nucleus received multiple inputs on the soma (upper arrows) and dendrites (lower arrows) from *Shox2+* neurons, indicating they are strongly driven by the *Shox2+* TCNs.

**Figure 1:**
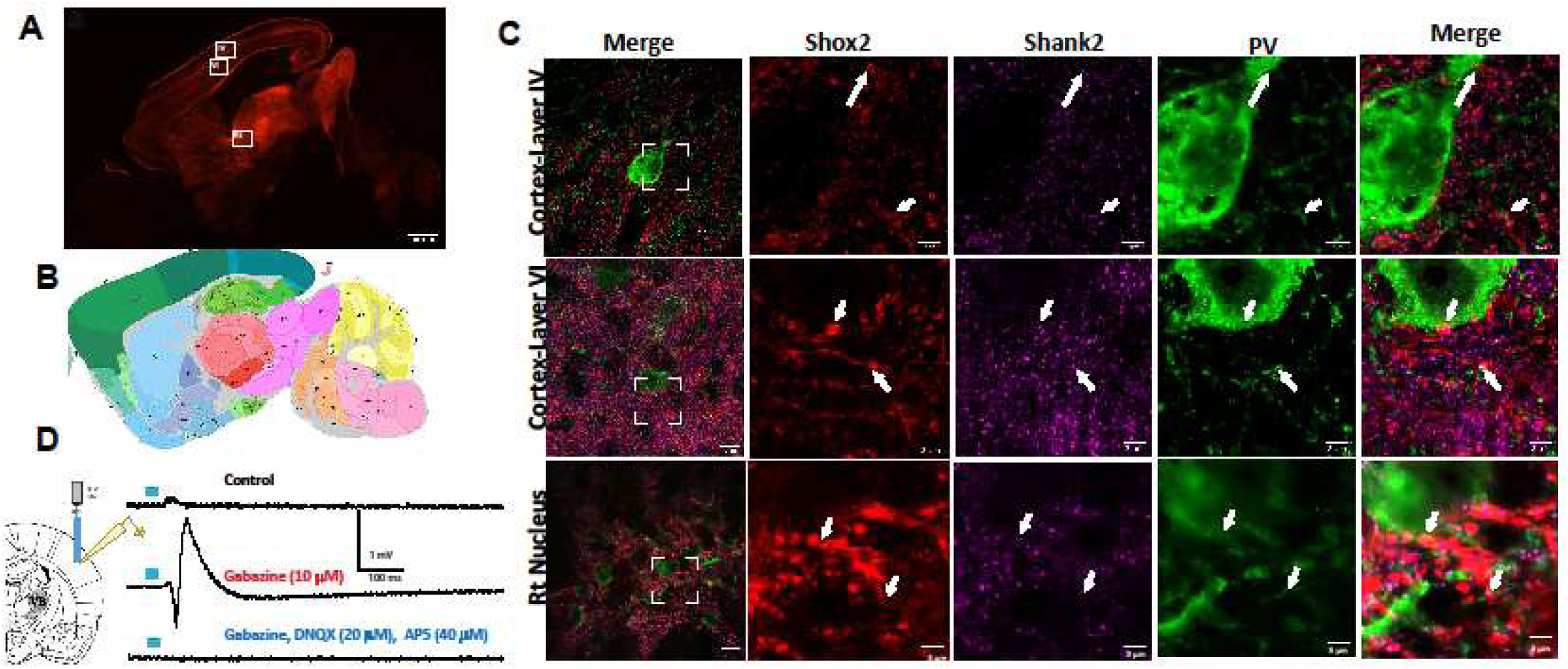
*Shox2+* TCNs possess typical TCN cortical and reticular connectivity properties. A: Sagittal image from a *Shox2*cre;Ai14tdT reporter mouse, with red fluorescent protein (RFP) as the reporter reveals localization of *Shox2* expression in the thalamus with projections to various layers of the cortex, including cortical layers IV (**box IV**) and VI (**box VI**), as well as projections to the reticular nucleus (**box Rt**). **B**: Allen Brain Atlas image of the same sagittal section. Abbreviations and link to sagittal image can be found <u>here</u>. **C:** Images of selected PV+ neurons within box IV, VI and Rt. Column 1: 100x images from boxed areas on sagittal image, from top to bottom: layer IV cortex, layer VI cortex, reticular nucleus (Rt). Column 2: Zoomed area of cell within the framed area from column 1 showing *Shox2*-expressing projections. Round ‘feet’ were determined to be presynaptic terminals. Column 3: Same area from column 1 showing Shank2 staining, puncta were determined to be glutamatergic synapse sites. Column 4: Same area of column 1 showing PV+ interneuron staining, areas with somas and dendrites were chosen. Column 4: Merged image. Areas of colocalization on cell somas (top white arrow for each row) and dendrites (bottom white arrow for each row) were identified as synapses. Arrows point to same location in rows 2-5. **D**: Thalamocortical slices were obtained from *Shox2*cre;ChRtdT mice. Field recording electrodes were placed within barrels of the layer IV barrel cortex (schematic on left). Selective activation of *Shox2*-expressing projections using 5ms blue light elicited an upward response (top trace/control). Application of 10µM gabazine to block GABAergic synapses reversed the polarity of the response and increased the amplitude (middle trace). Application of 20µM DNQX and 40µM AP5 to block glutamatergic synapses confirmed that the response was synaptic (bottom trace). Example traces, n=5.

To confirm functional connectivity to cortical cells, *Shox2+* TCNs were activated by crossing Shox2-Cre mice to a ChR2 reporter, and the response was recorded in layer IV of the cortex. Field potentials were recorded in the barrel cortex of thalamocortical orientation slices, a slice prep which preserves much of the VPM projection to barrel cortex (Agmon & Connors, 1991). Stimulation via a 5-ms blue LED light of the *Shox2+* cells in the VB resulted in a consistent cortical response (**figure 1D**). To dissect whether this response included inhibitory and/or excitatory elements, GABAergic synapses were blocked by bath application of 10µM gabazine. The same stimulus resulted in a response of increased amplitude—relative to control conditions—and of dual polarity. This indicates that *Shox2+* cells evoke a strong inhibitory response that suppresses excitatory responses within the barrels of the cortex. The remaining synaptic activity was completely blocked by AMPA and NMDA (glutamatergic) antagonists, 20µM DNQX and 40µM AP5, respectively. In the presence of these antagonists, the same stimulation failed to evoke a cortical response, confirming the previously recorded responses resulted from synaptic activity. These data indicate that *Shox2+* TCNs synapse in the layer IV barrel cortex and evoke excitatory and strong inhibitory responses. Collectively, these results demonstrate that *Shox2+* TCNs synapse on PV^+^ interneurons of the reticular nucleus, and interneurons and primary neurons of layers IV and VI of the cortex, which is a typical connectivity pattern of first order TCNs.

### Shox2 KD at P21 significantly affects burst firing and tonic firing of VB TCNs

We previously demonstrated that a global *Shox2* knockdown (KD) affects burst firing properties of TCNs of the anterior paraventricular thalamus. In thalamocortical circuitry, burst firing is elicited when TCNs are hyperpolarized by GABAergic reticular neurons (Steriade, Deschenes et al. 1985). This hyperpolarization is important as it affects two critical ion channels in the membranes of TCNs: hyperpolarization-activated cyclic-nucleotide-gated (HCN) channels and low threshold T-type calcium channels. Hyperpolarization activates HCN channels and deinactivates T-type channels. The deinactivation of the low threshold T-type channels allows them to be activated when membrane potentials are near resting membrane. The simultaneous opening of the HCN channels generates depolarization of the neurons to activate T-type Ca^2+^ channels, around -65mV. Activation of T-type Ca^2+^ channels generates a large, depolarizing, inward Ca^2+^ current. This current results in a wave of membrane depolarization that facilitates a cluster of action potentials—a burst. TCNs located in different thalamic nuclei exhibit varied burst firing properties (Desai and Varela 2021), thus, for a thorough investigation of Shox2 activity within the somatosensory system, confirmed that *Shox2* expression is also critical for VB TCNs firing properties.

To determine whether *Shox2* contributes to membrane and firing properties of VB TCNs, we generated a *Shox2* knockdown model by injecting an AAV2 virus, in which Cre is fused to GFP, into the VB of P21 *Shox2^fl/fl^*mice. At P28-35 patch-clamp recordings were obtained from GFP-expressing cells within the VB of these mice. Current-and voltage-clamp recordings were performed to isolate effects to TCN firing, membrane currents, resting membrane potential, and input resistance.

We tested whether *Shox2* expression is important for burst firing in VB TCNs by analyzing the rebound voltage response after hyperpolarization, or post-anodal burst, in control (*Shox2+*) versus KO VB TCNs. **Figure 2A** shows an example of control (left) and KO (right) TCN post-anodal bursts. Analysis of these bursts revealed that KO of *Shox2* at P21 significantly reduced the number of action potentials per burst (**figure 2B**) and area under the burst under the burst curve (**figure 2C**). In addition, the difference in voltage sag in response to hyperpolarization was measured (**figure 2D**). This sag response is attributed to activation of the HCN channels (Combe and Gasparini 2021), and was significantly reduced in the KO cells. In addition, KO of *Shox2* significantly increased input resistance (**figure 2E**), and significantly hyperpolarized the resting membrane potential (*Shox2+* -65.1 ± 1.5; *Shox2*KO -70.1 ± 1.1; p < 0.05; t = 2.1, df = 17), likely due to the reduced HCN current. Together, these results reveal that *Shox2* activity within VB TCNs affects membrane characteristics and is ultimately critical to their burst firing.

**Figure 2:**
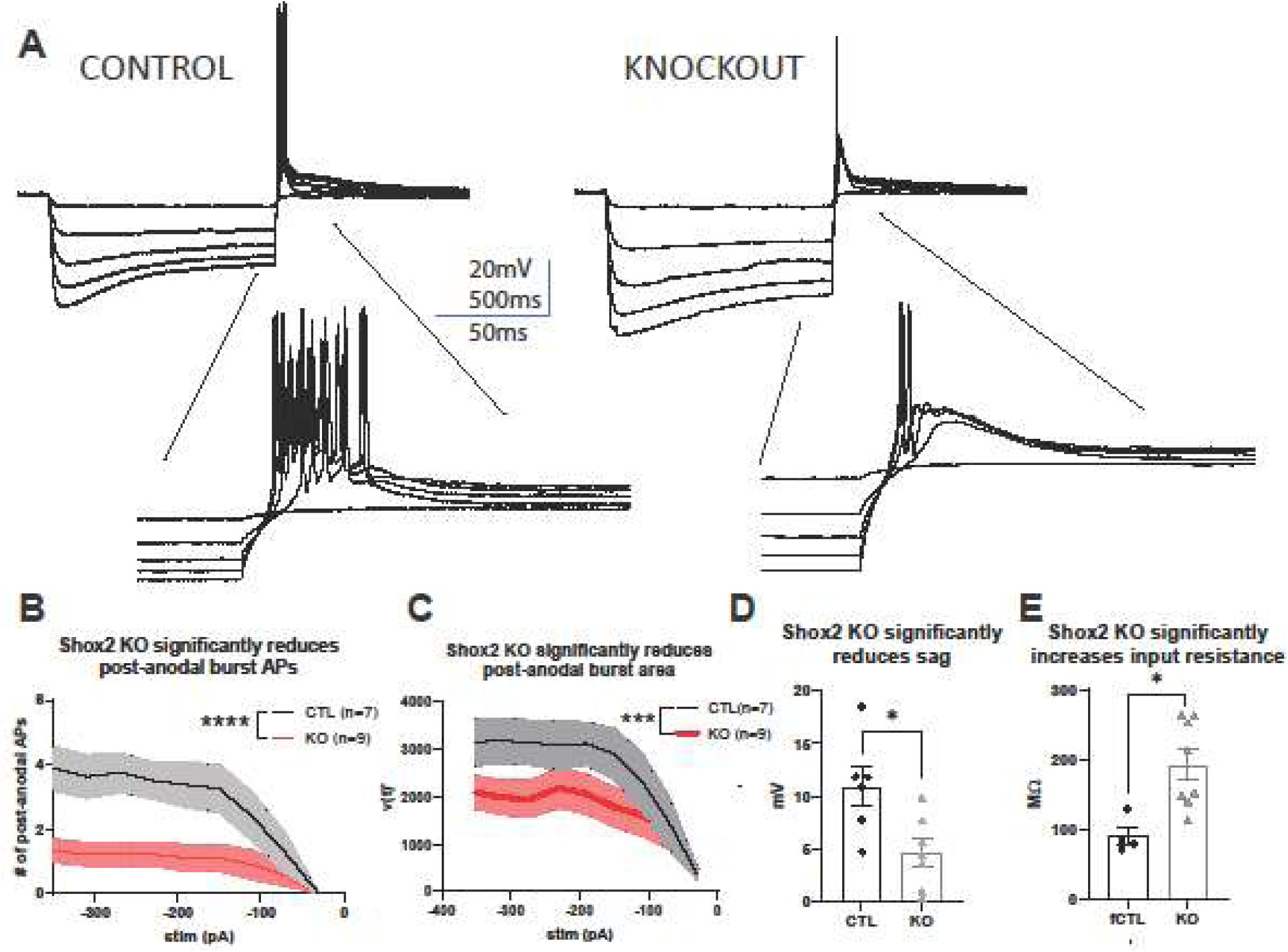
Shox2 is necessary for normal TCN burst firing. **A:** Example current-clamp recording from *Shox2*+ and KO neurons with a focus on post-anodal burst voltage response to various hyperpolarizing current injections. *Shox2*+ cells display robust burst responses with a cluster of multiple action potentials per burst. KO cell responses display a diminished low threshold spike and reduced number of action potentials per burst. **B:** *Shox2* KO significantly reduces number of post-anodal action potentials per burst. Black trace is the average of *Shox2+* TCNs, red trace is the average of *Shox2* KO TCNs. KO cells have significantly fewer action potentials per post-anodal burst (2-way repeated measures ANOVA, p<0.0001). **C:** *Shox2* KO significantly reduces area under burst curve. Area under burst curve was determined by taking the integral of the voltage response over time, upon release from varying hyperpolarizing current injections (-350pA in 40 pA steps). *Shox2* KO cells were determined to have a significantly smaller area under the burst curve as determined by 2-way ANOVA, p=0.0001). **D:** Quantification of sag reveals that *Shox2* KO significantly reduced HCN sag in response to hyperpolarization (student’s t-test, p=0.017). **E:** *Shox2* KO cells have significantly increased input resistance (student’s t-test, p=0.017).

Tonic firing of TCNs has also been shown to be critical to thalamocortical oscillations (Lee, Song et al. 2013, Amarillo, Zagha et al. 2014) and is tightly regulated by channels other than HCN and T-type (Kasten, Rudy et al. 2007). Tonic firing of VB TCNs in the S*hox2* KO cells was dysregulated. **Figure 3** shows representative voltage response traces of control *Shox2+* (**Figure 3B**) and KO (**Figure 3A**) TCNs in response to a range of depolarizations. Analysis of all tonic firing revealed that depolarization of *Shox2+* TCNs had an average rheobase of 50 pA and saturated at an average firing rate of 50 Hz with an injection of 400 pA of current (**Figure 3C**). However, KO TCNs had a much higher rheobase of over 100 pA, and at 400 pA of current injection they averaged 17 Hz (**figure 3C**). Further analysis showed that the peak amplitude of action potentials during tonic firing was significantly reduced in KO cells (**figure 3D**), but there was no effect on action potential halfwidth (p = 0.26; **figure 3E**) or time to action potential peak (p = 0.11; **figure 3F**). These results show that *Shox2* KO caused aberrant burst and tonic firing in VB TCNs and altered membrane properties.

**Figure 3.**
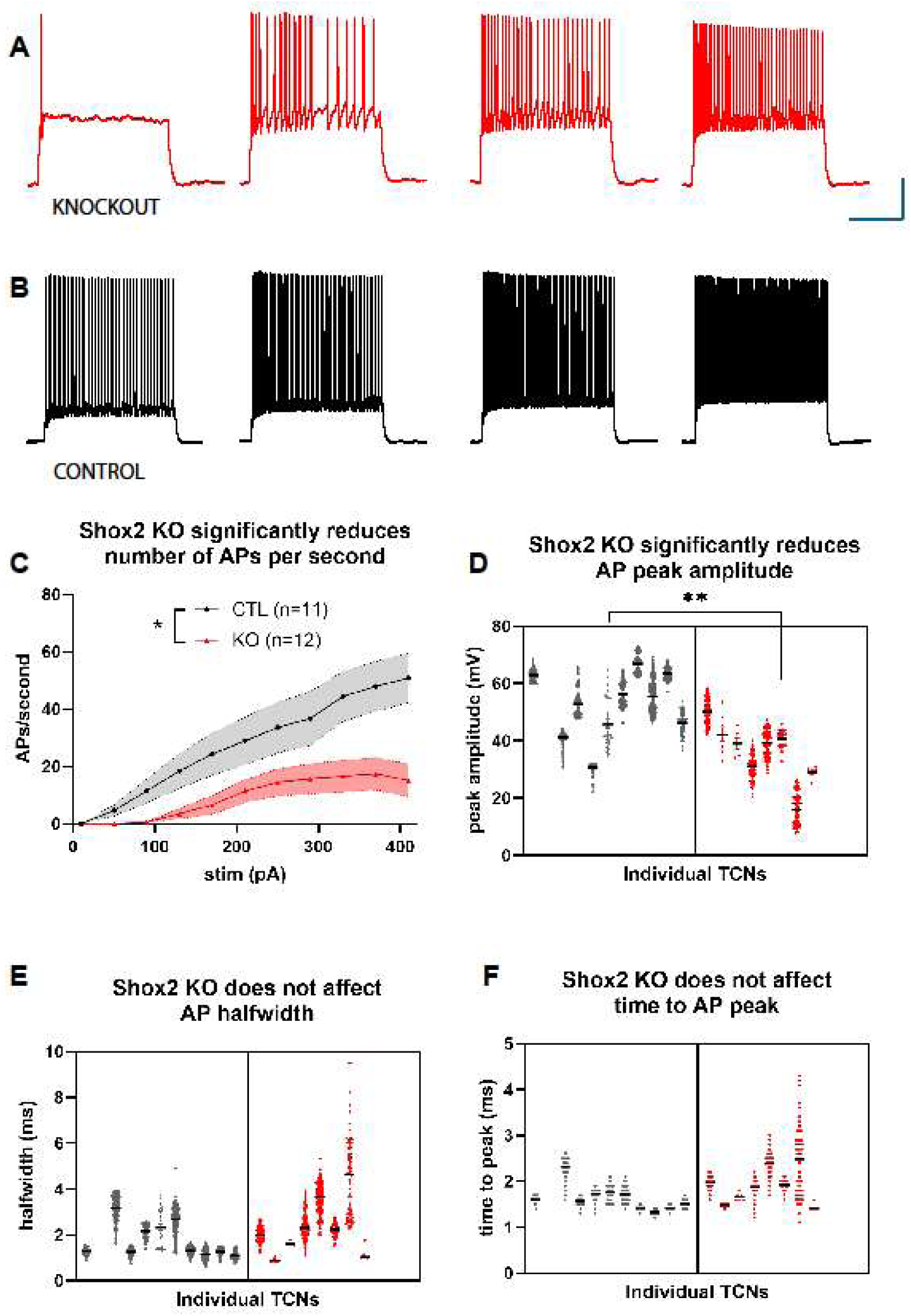
Shox2 expression is necessary for normal tonic firing of VB TCNs. **A**: Representative traces of *Shox2KO* and **B:** *Shox2+* TCNs in response to a 110-350 pA injection of current from a membrane potential of –60 mV**. C:** Saturation curves of action potentials (APs) per second in CTL (black) and KO cells (red) in response to increasing current injections. KO cells have significantly fewer APs/s (2-way ANOVA, p=0.013**). D:** APs in KO cells are significantly smaller in amplitude (nested t-test, p=0.006). **E:** KO does not significantly affect AP halfwidth. **F:** KO has no effect on time to AP peak.

*Shox2* KD does not affect gross cortical target organization but does impair somatosensory perception Since *Shox2* KO caused a significant disruption of firing properties during burst and tonic firing in *Shox2* KO TCNs, we further investigated the consequences of this disruption of firing properties in the cortex. The cortex receives these burst and tonic firing signals from TCNs during development and adulthood, and cortical receipt of these signals influences the formation and function of the thalamocortical circuit. Elimination of thalamic input to the neocortex suggests that thalamic input is not necessary for regionalization of gene expression in the cortex (Miyashita-Lin, Hevner et al. 1999) but is important for refinement of cortical circuitry. For example, in the absence of Vglut2 in TCN terminals which prevents glutamatergic transmission, gross cortical barrel formation is abolished (Li, Fertuzinhos et al. 2013).

Given *Shox2*’s effect on thalamic activity above and its role in development, gross cortical barrel formation was investigated. The critical window for barrel formation is near P7 (Erzurumlu and Gaspar 2012), and, thus, we hypothesized that loss of VB *Shox2* at P6, but not P21, would disrupt barrel formation. To test this hypothesis, unilateral VB *Shox2* knockout models were made at P6 and P21. This model allowed for a within mouse control in which we compared cortical barrels ipsilateral to the injection versus barrels contralateral to the injection of the same mouse (**schematic: figure 4A**). At PND 35 left and right thalami were processed for *Shox2* mRNA to confirm KO (**figure 4B**), and successful knockdown at both ages was confirmed. Barrel formations were visualized in flattened slices of the barrel cortex stained with cytochrome oxidase (CO) in the KO and control sides (**figure 4D**). There was no effect on the barrel map, since all are present in both the P6 and P21 models. The absence of effect in our P6 KO model could be because the virus may take multiple days to fully affect a thalamic nucleus, thus, by injecting at P6—only one day before the critical development window—we may have knocked down *Shox2* after the barrels were formed. To address this, we repeated our experiment with P3 unilateral, ultrasound injections. However, analysis reveals that this timepoint injection still fails to disrupt gross barrel formation (data not shown).

**Figure 4:**
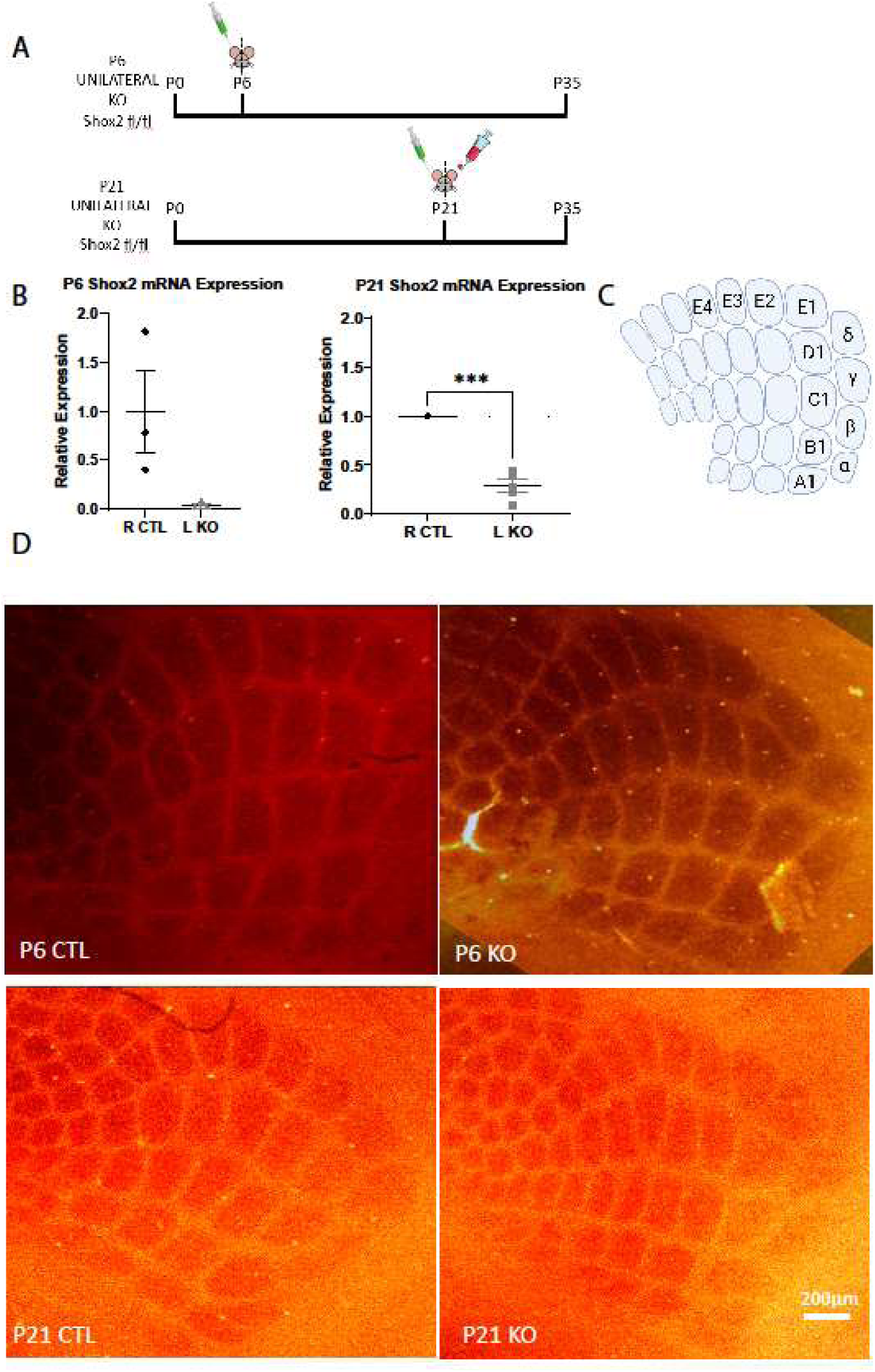
Shox2 is not important for cortical target organization. **A:** Schematic diagram of unilateral P6 and P21 KO mouse experimental timeline. Top: P6 *Shox2*^fl/fl^ mice received unilateral VB injections of an AAV2-GFPcre virus ultrasonically. Bottom: P21 *Shox2*^fl/fl^ mice received unilateral, VB injections of an AAV2-GFPcre virus and an RFP control virus stereotactically. At P35, cortex from both sets of mice were analyzed for *Shox2* mRNA expression in the VB. **B:** *Shox2* mRNA expression from VB punches confirms a reduction of *Shox2* mRNA in the KO versus CTL side in both P6 and P21 models. **C:** Flattened view of barrels, map of flattened barrel cortex where Greek letters and alpha-numeric numbering outlines the border of the full barrel cortex. **D:** CO staining was performed on flattened P35 barrel cortices of unilateral VB KO at either P21 or P6. Sections were imaged and images were aligned to generate a reconstruction of barrel fields from P6 (top row), P21 (middle row) control and KO sides. Upon comparison to the barrel cortex map, all barrels are present at each KO timepoint, thus no gross differences were observed.

### Sensorimotor disruption

Although gross disruption of cortical barrel development was not observed, we questioned whether the observed disruption of normal firing activity of TCNs was sufficient to impair sensorimotor function. To test this, sensorimotor function in a global *Shox2* KO mouse as well as the VB KD model was tested. KO and control mice were tested on the adhesive removal tests (Bouet, Boulouard et al. 2009, Freret, Bouet et al. 2009). When a small piece of adhesive tape was placed on the foot of the global KO mouse, there was a significantly increased latency to removal, indicating that *Shox2* KO mice have deficits in sensorimotor function (**figure 5A**). To test if this effect was specifically localized to the VB nucleus of the thalamus, sensorimotor function in the VB KD was tested.

**Figure 5:**
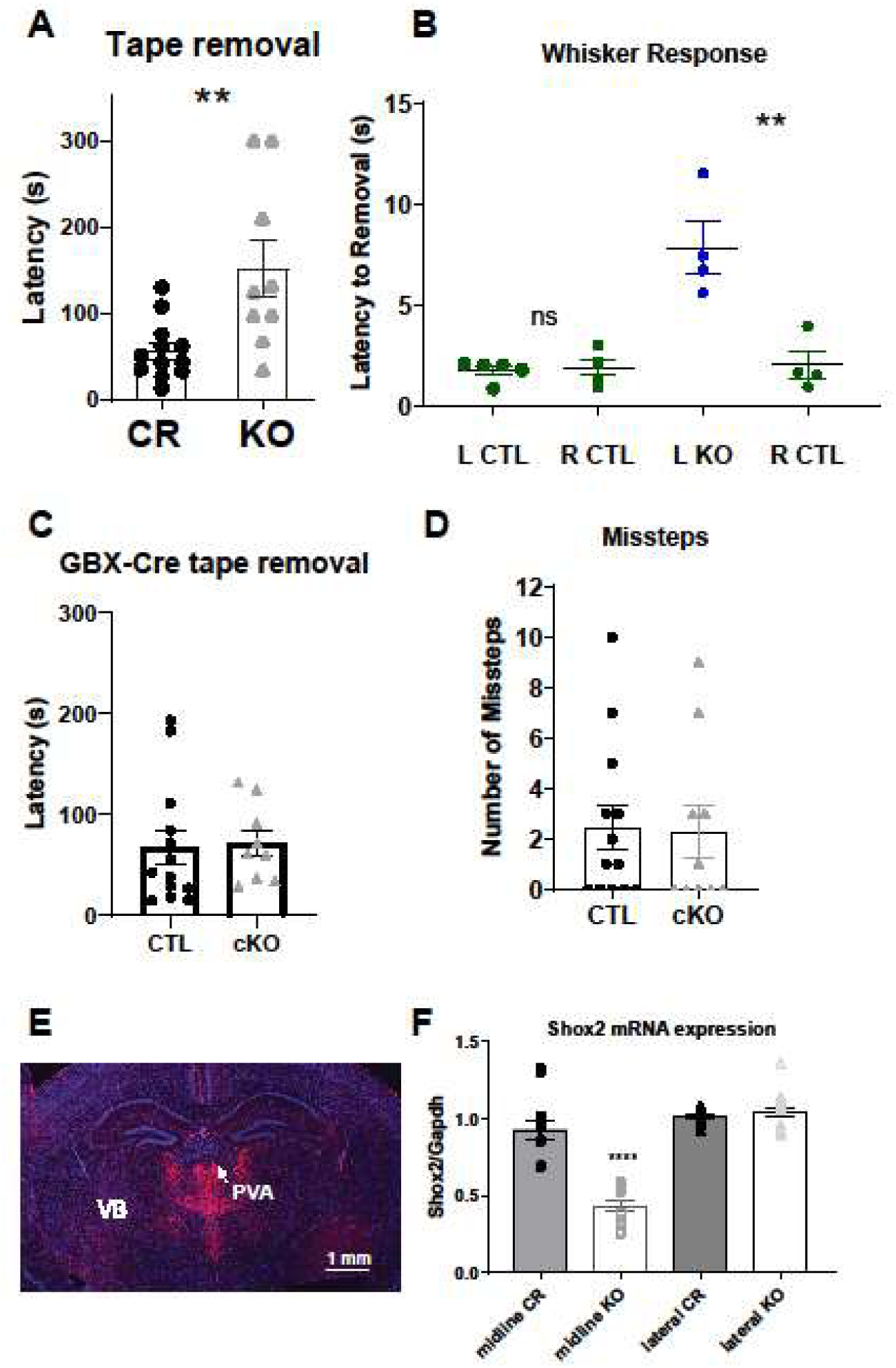
*Shox2* expression is important for somatomotor function. **A**: Increased latency to removal of adhesive tape placed on the foot in a tamoxifen-inducible, global *Shox2* KO mouse versus littermate controls, indicating that *Shox2* KO mice have a deficit in somatomotor function. **B:** In a targeted VB, unilateral *Shox2* KO mouse (*Shox2^f^*^l/fl^ and stereotactically injected cre virus), stickers were placed on either left-or right-side whiskers and latency to removal was recorded. Each value is an average of three trials. **Left:** Control (CTL) subjects are non-injected Shox2^fl/fl^ mice to test for handedness. There is no significant difference between left-and right-side latency to removal, which indicates that there is no pre-existing handedness. **Right:** A within mouse study in which *Shox2* was knocked down in the right VB, corresponding with contralateral left whiskers (L KO), and the left control VB was injected with a control virus (R CTL). KO of *Shox2* in the VB nucleus of the thalamus significantly increases latency to tape removal. **C:** Midline-thalamus KO of *Shox2* does not impair somatosensation: There is no significant difference in the latency to removal of sticky tape from a hind paw as determined by a student’s t-test. **D:** By analysis via the grid walking test and number of missteps, Shox2 KO mice exhibited no gross motor function deficits. **E:** GBXCre;*Shox2*^fl/fl^, midline-thalamus KO, Cre expression in the midline of the thalamus in red and **F:** *Shox2* mRNA expression confirming KD of *Shox2* transcription in the midline and not the lateral/sensory thalamus.

The adhesive removal sensory test was adapted by applying the sticker to the whiskers instead of the paw, since the VB nucleus processes sensory information from the whiskers (Jones 2002). Stickers were placed on either left-or right-side whiskers and latency to removal was recorded. In addition to the within mouse control, the non-injected Shox2^fl/fl^ mice were tested for handedness. There was no significant difference between left-and right-side latency to removal, which indicates that there is no pre-existing handedness relevant to this task (**figure 5B**, **left**). *Shox2* was knocked down in the right VB, corresponding with contralateral left whiskers (L KO), and the left control VB was injected with a control virus and corresponds with the right whiskers (R CTL). Time to removal of the sticker from the left whiskers was significantly increased on the left side compared to the right, indicating that KD of *Shox2* in the VB nucleus of the thalamus significantly impairs sensorimotor through the whiskers (**figure 5B**, **right**). These data indicate that *Shox2* KD in the VB is sufficient to impair somatosensory perception through the whiskers, possibly via the mechanism of reduced TCN action potential density.

We also tested sensorimotor function using the GBX driver to KO *Shox2* (GBX-Cre,*Shox2^fl/fl^*). In these animals, we do not expect *Shox2* KO in the VB as at the age of tamoxifen delivery, GBX is primarily expressed only in midline nuclei, and we confirmed that this transgenic cross knocked down expression of *Shox2* mRNA in the midline, but not lateral thalamic nuclei (**figure 5E,F**). No change in latency to removal was observed between control and KO mice. To determine if these mice exhibited any effects on gross motor function, we also assessed gross motor function via the grid walking test in the GBX-cre *Shox2* KO mice. No significant effects were observed on time across the grid (KO: 29.76± 17.1s, n = 10; Ctrl: 19.72± 4.7s, n = 13; p = 0.54), missteps (**Figure 5D**; 2.3 ± 1 control – 2.46 ±0.9; p = 0.9) or full falls (0.5 ± 0.4, ctrl 0.15 ± 0.1). This suggests that motor function is not disrupted in the GBX-cre mice.

### *Shox2* KO at p21 significantly reduces spindle density

Proper firing of TCNs is critical for the generation of sleep spindles within the thalamus (Swadlow and Gusev 2001, Fernandez and Luthi 2020). In mice lacking TCN T-type Ca channels, TCNs lack the ability to burst fire, and in mice lacking the α1_G_-subunit of TCN T-type Ca^2+^ channel, preventing current through the channel, sleep spindle generation is disrupted (Lee, Kim et al. 2004, Crunelli, Cope et al. 2006, Astori, Wimmer et al. 2011, Pellegrini, Lecci et al. 2016, Fernandez, Vantomme et al. 2018) . Further, HCN4 deficient mice have significantly fewer evoked TCN bursts and slower cortical oscillations (Zobeiri, Chaudhary et al. 2017, Zobeiri, Chaudhary et al. 2019). Tonic firing of TCNs has also been proposed to be critical for spindle generation (Lee, Song et al. 2013). Thus, we hypothesized that the significant reduction in burst and tonic action potentials observed in VB of *Shox2* KO TCNs would disrupt thalamus generated oscillations, specifically sleep spindles. A previous study demonstrated that spindle rate during non-REM sleep increases significantly before the onset of rapid eye movement sleep but not before wakefulness (Bandarabadi, Herrera et al. 2020), suggesting a role for spindles in deepening sleep. Therefore, we determined total spindle frequency during NREM and the last minute leading up to REM and wake.

To determine whether Shox2 expression in the thalamus is important for sleep spindles generated by the thalamocortical circuit, EEGs from the previously described targeted KO of VB *Shox2* were measured and compared their cortical activity to littermate controls. We then recorded cortical EEG and EMG activity in 8 control and 8 KO mice over 24 hours (**Figure 6A**). Duration of sleep and cortical EEG power and spindle activity were determined (**Figure 6B**). No differences in duration or rhythms of sleep—in either males or females (**Figure 6C**)—nor was there a difference in power across a spectrum of frequencies. Reduced sleep spindle density, defined as number of spindle events per minute, in KO mice during NREM sleep was observed (**Figure 6D**). We tested total spindle rate and spindle rate in the minute leading to REM sleep and wake (**Figure 6E**). We found a significant effect of genotype and phase and an interaction of genotype and phase, indicating that spindle rate decreased in KO mice and the KO mice showed no increase prior to REM sleep as has previously been reported in WT mice (Bandarabadi, Herrera et al. 2020),.

**Figure 6:**
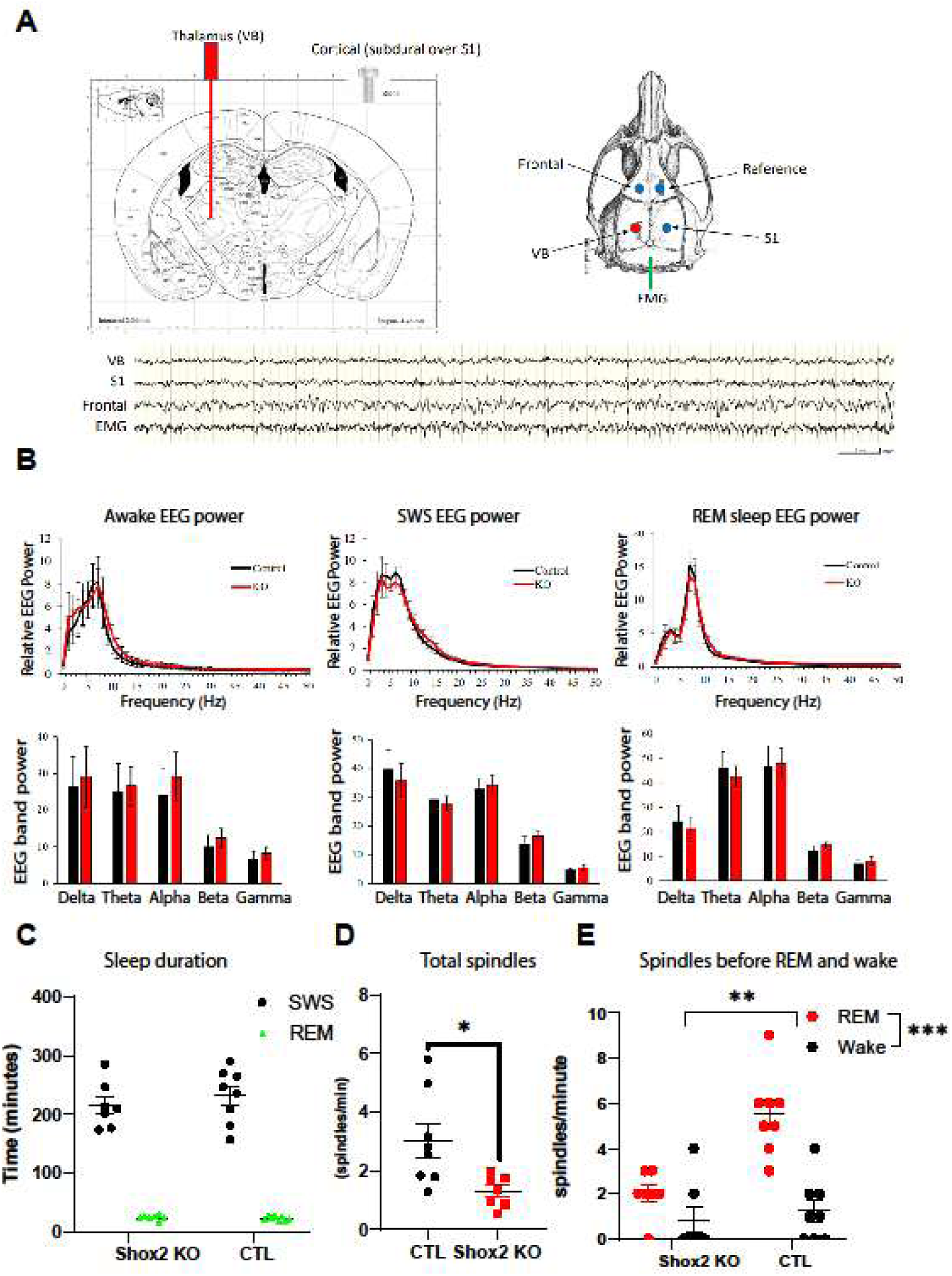
*Shox2* expression is important for thalamocortical spindle density. **A:** Top: Diagram showing coronal location of VB probe placement (left) and dorsal location of frontal, reference, VB, S1, and EMG probes (right). B. Power analysis of cortical EEGs reveals no differences in oscillation frequency power between control or KO mice during awake (left), slow wave sleep (middle) or REM (right). Bottom: Example trace recordings from each probe. **B:** Frequency power graph (above) and power of various rhythms (below) from awake (left), slow wave sleep (middle) and REM (right). Below – power of various rhythms during awake **C:** Time spent in slow wave sleep (SWS) and REM sleep is not different between control and KO mice. **D:** *S*WS spindle density is significantly reduced in *Shox2* KO mice. A bandpass filter of 9-16Hz was used to analyze the frontal EEG of CTL v KO mice during SWS. Analysis reveals that Shox2 KO mice have a significantly reduced density of sleep spindles (spindles/min) as determined by a student’s t-test, p=0.017. **E**. Spindle density in the minute leading up to REM and wake was determined. While the CTL mice showed a significant increase in spindle frequency leading up to REM, the KO mice did not.

### Shox2 KO significantly reduces memory formation

Sleep spindles are functionally associated with memory consolidation during sleep; increased spindle activity enhances memory consolidation, and spindle disruption impairs memory consolidation (Latchoumane, Ngo et al. 2017). Thus, we hypothesized that the reduced sleep spindle density in Shox2 KD mice would translate to a deficit in memory consolidation.

### Memory consolidation

In the global *Shox2* KD mice, an open field test was used to determine that total activity was not altered (**figure 7A**) and, interestingly, anxiety levels were decreased in KD mice as measured by percent time spent in the center of the open field (**figure 7B**). The novel object recognition task was used to test memory consolidation in the global *Shox2* KD mice (**figure 7C**). KD and littermate control mice were familiarized with two of the same object for 5 minutes and the discrimination index (described in methods), based on interaction time, between the two groups was quantified and no differences were observed (**figure 7D**, **left**). Mice were returned to their home cages and, 24 hours later, when one of the objects was replaced with a novel object of similar size and texture, mice were allowed to interact with either object for 5 minutes (**figure 7D**, **right**). *Shox2* KO mice showed a significantly decreased discrimination index between the two objects, indicating a deficit in novel object recognition.

**Figure 7:**
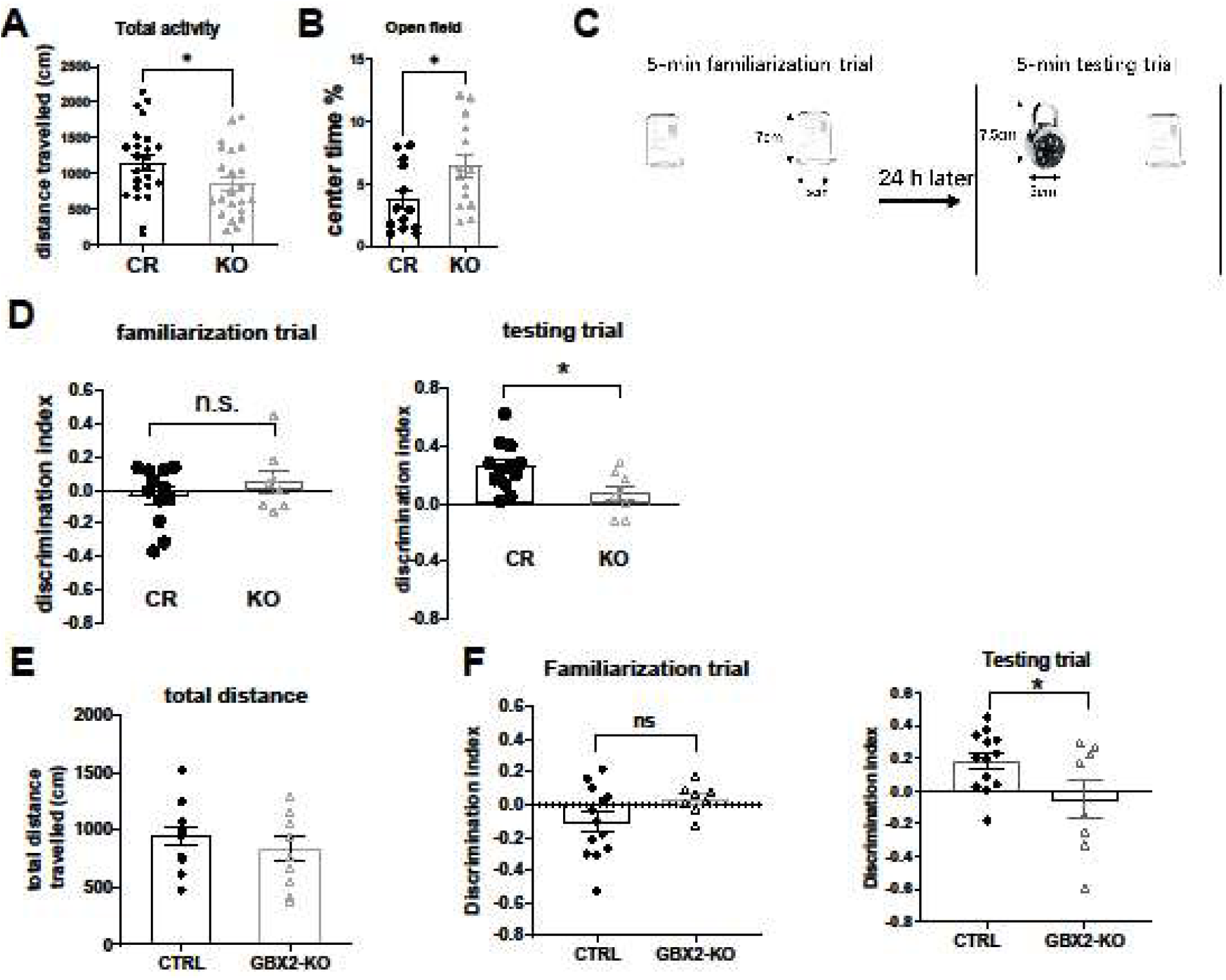
*Shox2* expression is important for memory consolidation. **A/B:** Tamoxifen-inducible, global KO mouse model was tested on total activity and open field test. *Shox2* KO showed reduced distance **(A)** but increased time in center of open field **(B)** suggesting less anxiety (*, p<0.05). **C:** Schematic of novel object recognition. **D**: Mice were familiarized with two of the same object for 5 minutes. The discrimination index, based on interaction time, between the two groups was quantified. No difference between control and KO mice was observed on the familiarization trial(D, left). Mice were returned to their home cages, and, 24 hours later, when one of the objects was replaced with a novel object of similar size and texture, mice were allowed to interact for 5 minutes. (D, right): During the testing trial, *Shox2* KO mice showed a significantly decreased discrimination index, indicating a deficit in novel object recognition. **E.** *Gbx2Cre:Shox2* KO mice were tested on exploration and novel object recognition. No differences in total distance were observed. **F:** *Gbx2Cre:Shox2* KO mice and littermate controls showed no significant differences in object investigation during the familiarization trial (left). During the testing trial, KD mice showed a significantly decreased discrimination index, indicating a deficit in novel object recognition (right).

Since a sensory perception impairment in the global *Shox2* KD mice, it is possible that this impairment of sensory perception was a confounding variable in the novel object recognition task. To rule out this possibility, the same novel object recognition task in a mouse model where *Shox2* was selectively knocked down along the midline of the thalamus (memory associated nuclei), but still maintained expression in the sensory nuclei—our GBXCre;Shox2^fl/fl^ mice as discussed above that did not exhibit a somatomotor effect. There was no significant difference in total distance explored in these mice (F**igure 7E**) However, when tested on novel object recognition, the midline-thalamus, *Shox2* KD mice still presented with a significantly reduced discrimination index, indicating non-sensory reliant, memory consolidation impairment (**figure 7F**).

## Discussion

Because of *Shox2’s* potential role in neurodevelopmental disorders and high levels of *Shox2* expression in thalamocortical neurons, we investigated it as a candidate transcription factor to coordinate thalamocortical neuron functional development and structural connectivity. We previously showed that *Shox2* is expressed in TCNs during development and into adulthood, and conditional knockout of *Shox2* in the young adult regulates expression of genes important for TCN cell structure and function. Furthermore, lack of *Shox2* affects TCN cell excitability which leads to increased seizure susceptibility in *Shox2* KD mice (Yu, Febbo et al. 2021). Building on our previous work, in the current study we demonstrate the critical role of *Shox2* expression in the development and function of thalamocortical neurons (TCNs) of the VB nucleus. Specifically, we show that *Shox2* is expressed in thalamic neurons that synapse on cortical and thalamic reticular interneurons. *Shox2’*s expression is crucial for TCN function and robust tonic and bursting properties of TCNs. These firing patterns contribute to the thalamus’ role in somatosensory function as well as sleep spindles, that contribute to memory formation and have implications for neurodevelopmental disorders such as autism and schizophrenia.

While no syndromes have been directly linked to SHOX2 mutations in humans, this work has clinical implications, as the human *SHOX* family contains *SHOX* and *SHOX2*. *SHOX* is a paired-related homeobox gene and encodes a transcription factor involved in cell cycle and growth (Rao, Blaschke et al. 2001). The *SHOX* locus is within the pseudoautosomal region 1 of the sex chromosomes, and escapes X inactivation (Blaschke and Rappold 2006). Heterozygous mutations or deletions of *SHOX* are also associated growth disorders, including Leri-Weill dyschrondrosteosis and Turner Syndrome (Belin, Cusin et al. 1998, Blaschke, Monaghan et al. 1998, Cormier-Daire, Belin et al. 1999, Blaschke and Rappold 2006, Blaschke, Hahurij et al. 2007, De Sanctis, Tosetto et al. 2012, Choi, Seo et al. 2015, Benito-Sanz, Belinchon-Martinez et al. 2017). Further, microduplications of *SHOX* affecting the 3’-enhancers are enriched in autism spectrum disorders (Tropeano, Howley et al. 2016), and duplications affecting the 5’ enhancers are typically seen in patients with skeletal dysmorphism *SHOX* (Durand, Roeth et al. 2011). Further, patients with these syndromes are at increased risk for various neurodevelopmental pathologies, including learning disabilities and developmental delay, epilepsy, and abnormal visuospatial, social, and executive function.

Although these neurodevelopmental deficits are consistent with disrupted thalamic function, the mechanisms of neural impairments associated with SHOX dysregulation are unknown. Disruption of TCN firing properties, similar to what we have observed from *Shox2* KO, likely underlies some neurodevelopmental disorders. The rodent genome does not possess the SHOX gene, but *Shox2* is a closely-related paralog with identical DNA-binding homeodomains. Prior evidence supports that mouse *Shox2* is closely related to h*SHOX* and *hSHOX2* (Rosin, Abassah-Oppong et al. 2013), and importantly, human SHOX can functionally replace *mouse* Shox2 in regulating embryonic development and postnatal survival (Liu, Chen et al. 2011). Furthermore, *Shox2* is directly repressed by Aristaless-related homeobox *Arx*, a gene implicated in intellectual disability, seizures, and autism (Fulp, Cho et al. 2008, Colasante, Sessa et al. 2009), and *Shox2* is significantly up-regulated in an *Arx* mouse model of infantile spasms (Olivetti, Maheshwari et al. 2014). These results suggest that *Shox2* may also play a role in these disorders downstream of *Arx*. Understanding the contribution of *Shox2* and downstream effectors to the molecular function of TCN neurons and the thalamocortical interactions of *Shox2-*expressing neurons is therefore critical to obtain an overall understanding of the role of thalamocortical neurons in cognitive function in general.

### Regulation of transcriptional control in the thalamus

How *Shox2* regulates mechanisms important for neuronal function in the thalamus is unknown. Pioneer factors are transcription factors that can modulate transcription by directly binding to condensed chromatin and can interact with other transcription factors or histone modifying enzymes. These factors are commonly mutated or mis-regulated in cancer and are key transcription factors to generate induced pluripotent stem cells. Several important pieces of evidence suggest that *Shox2* may act as a pioneer factor in other tissues. First, SHOX2 promotes cell division and is frequently hypermethylated in lung cancer tissue (Kneip, Schmidt et al. 2011, Darwiche, Zarogoulidis et al. 2013, Zhao, Guo et al. 2015, Zhang, Yu et al. 2017). In addition, a previous study demonstrated that SHOX2 overexpression in embryonic stem cells upregulates a cardiac pacemaker gene program that results in enhanced automaticity (Ionta, Liang et al. 2015), suggesting that SHOX2 is a master regulator of biological pacing. Furthermore, previous studies have demonstrated drastically reduced accessibility of chromatin regions in *Shox2* KO cells (Xu, Wang et al. 2019). *Shox2* binding sites across different tissues by ChIP-Seq and identified enrichment of three distinct motifs: the Meis/Pbx motifs in the Shox2 palate binding sites, the Hox9-TALE related motifs in the Shox2 limb binding sites (Ye, Song et al. 2016), and the Nkx2-5 binding motifs in the heart (Ye, Wang et al. 2015). These differential binding motifs indicate tissue-specific DNA binding sites of Shox2 results from its physical interactions with other transcription factors whose conserved motifs are present in the Shox2 binding sites. Future work will address the mechanisms through which Shox2 regulates gene expression in the thalamus and if Shox2 acts as a pioneer factor that contributes to an upregulation of a thalamocortical neuron program.

### Transcriptional regulation of ion channels

Previous studies indicate that combinatorial regulation by transcription factors may be important for control of ion channel gene expression (Wolfram, Southall et al. 2014). In the adult mouse thalamus, combinatorial expression of Tcf7l2, Lef1, Gbx2, Prox1, Pou4f1, Esrrg, and Six3 define molecular divisions that loosely correlate with established nuclei divisions (Nagalski, Puelles et al. 2016). Pax6 and Gli2 have been identified as transcription factors that are critical to the development of thalamocortical projections (Pratt, Vitalis et al. 2000, Callejas-Marin, Moreno-Bravo et al. 2022).TCF7L2, expressed throughout the thalamus as well as the habenula, has also been shown to be critical for thalamocortical projections and habenula organization (Bem, Brozko et al. 2019). Few studies, however, have addressed the maintenance of TCN firing properties in developed thalamus. Our investigations are the first to demonstrate that Shox2 is important for maintenance of the TCN bursting program. Here, we show that these properties are important for spindle oscillations, sensorimotor function and object memory. ^i^

### Spike timing and sensory perception

Here, we have shown Shox2 expression is important for maintenance of VB-TCN spike timing. It is well established that thalamic gating of cortical sensory information is spike-timing dependent (Sherman, 2001a; Sherman, 2001b; Swadlow and Gusev, 2001; Swadlow, 2002; Lesica and Stanley, 2004; Lesica et al., 2006; Wang et al., 2010; Stanley et al., 2012; Whitmire et al., 2016, Borden et al 2022). While spike timing and synchrony promote sensory processing, disruption of timing impairs it (kim & shin et al 2003, Sukchan et al 2008, Simon and Wallace 2016). These studies, along with the absence of sensorimotor deficits in the GBXcre-Shox2 KO mice, support our conclusion that Shox2 KO deficits in the somatomotor test performance derived from an impairment of sensory perception, rather than motor deficiencies. Further, although we focus on the somatosensory circuit in this study, *Shox2* is expressed throughout the thalamus, with a cortical target pattern that is typical of first order TCNs. This suggests that the activity of *Shox2-*expressing TCNs is involved in the processing of primary, thalamic sensory information.

### Sleep spindles and memory consolidation deficits

Sleep spindles are generated in the thalamus through reciprocal connections between the dorsal thalamus and the reticular nucleus. Functionally, sleep spindles have been associated with memory consolidation during sleep in that increased spindle activity enhances memory consolidation, and spindle disruption impairs memory consolidation (Latchoumane et al., 2017). Furthermore, the targeted KO mice did not show an increase in spindle frequency just prior to REM as was shown previously (Bandarabadi, Herrera et al. 2020) and in our control animals. This increase in sleep spindle frequency is thought to be important for deep sleep, but the functional significance of this effect needs further investigation.

Memory consolidation relies on the integration of multi-regional brain activity, including the thalamus (Shaw and Aggleton 1995) and specifically the mediodorsal nucleus (Chao Nickolaus Wang Huston 2022). We showed significantly impaired object recognition in 2 different S*hox2* KO lines (global and GBxcre). It is possible that a sensory deficit could contribute to the object recognition deficit, however, importantly the GBXcre mice, which did not show any sensorimotor impairments also showed significantly impaired object location memory. The reduced sleep spindle density was recorded in the VB KD mouse, a nucleus whose TCN burst, and tonic firing has been directly shown to generate cortical sleep spindles (Urbain, 2024). However, from our experiments we do not conclude that spindle activity generated in primary nuclei affect memory, since the deficits in memory consolidation were observed in the global Shox2 KO. In future studies, it would be of interest to investigate memory consolidation in Shox2-VB KD to address this. Regardless, the fact that we observed a significant cortical effect on spindles when knocking down Shox2 in a single thalamic nucleus speaks to the critical importance of thalamic Shox2 expression in the generation of cortical spindles. This is notable because reduced sleep spindle density has been identified as a biomarker for autism and schizophrenia (Copping & Silverman, 2021; Manoach et al., 2016).

### Conclusion

In conclusion, we present Shox2 as a protein involved in proper thalamocortical circuit function by showing that its expression within TCNs of the VB is necessary for TCN firing properties, including burst and tonic firing. Expression of Shox2 promotes spindle oscillations, sensory perception, and memory consolidation in mice. This implicates Shox2 as a possible molecular target for thalamic dysfunction and its behavioral correlates, such as autism and schizophrenia.

## Methods

### Mice

All animal procedures were approved by Tulane University Institutional Animal Care and Use Committee (IACUC) according to National Institutes of Health (NIH) guidelines. *Shox2* transgenic mice including *Shox2^Cre^*, *Shox2^LacZ^*, *Shox2^f/f^* and *Rosa26^CreERt^*mice were generously donated by Dr. Yiping Chen. All wildtype C57Bl/6N mice were ordered from Charles River. *Rosa26^LacZ/+^ (stock #003474) and Gbx2^CreERt/+^* breeders (stock #022135) B6.Cg-Gt(ROSA)26Sortm27.1(CAG-COP4*H134R/tdTomato)Hze/J https://www.jax.org/strain/012567 were ordered from Jackson Lab.

In inducible KO experiments, *Rosa26^CreERt/+^,Shox2^f/f^* or *Rosa26^CreERt/CreERt^,Shox2^f/f^* female mice were crossed with *Shox2^-/+^*male mice or, in the case of the *Gbx2* animals, *Gbx2^CreERt/+^*,*Shox^f/f^*or *Gbx2^CreErt/CreErt^,Shox^f/f^* were crossed with *Shox2*^-/+^ male mice. Litters were labelled and genotyped at postnatal day 10. The KO group was the *Rosa26^CreERt/+^*, *Shox2^-/f^* mice (*Gbx2^CreRt/+^, Shox2^-/f^*) and the control (CR) group was the littermate *Rosa26^CreERt/+^ (Gbx2^CreErt/+^),Shox2^+/f^*. In the *Shox2^LacZ/+^* and *Shox2^Cre/+^* mice, the first two exons of the *Shox2* allele were partially replaced by *LacZ* and *Cre* genes respectively in order to obtain the expression of *LacZ* mRNA and *Cre* mRNA under the control of the *Shox2* promoter, while the unaffected alleles express *Shox2* mRNA (Sun, Zhang et al. 2013). The *Rosa26^CreERt/+^* mouse line is a transgenic mouse line with a tamoxifen-inducible Cre^ERt^ inserted in the *Rosa26* loci. The *Rosa26^LacZ/+^* mouse line is a transgenic mouse line with a floxed stop signals followed by LacZ gene inserted in the *Rosa26* loci (Soriano 1999). This ‘cre reporter’ mouse strain was used to test the expression of the *Cre* transgene under the regulation of a specific promoter.

The Rosa26^CreErt^ is a global KO, whereas the *Gbx2^CreERt^*mouse was used to knockdown *Shox2* specifically in the medial thalamus. Our localization studies with *Gbx2*-promotor driven GFP staining showed that *Gbx2* promotor-driven *CreERt* is specifically expressed in the midline thalamus of the Gbx2^CreErt^ adult mouse (Fig. 5E). Further testing using RT-qPCR showed that *Shox2* mRNA was reduced in the medial thalamus of the *Gbx2^CreRt/+^, Shox2^-/f^* compared to CR mice, but not lateral thalamus (Fig. 5F).

To induce recombination in animals bearing a Cre^ERt^, pre-warmed tamoxifen (100-160 mg/kg) was injected intraperitoneally into KO mice and CR littermates at the same time every day at PND 21 for five consecutive days. Tamoxifen (20 mg/mL) was dissolved in sterile corn oil (Sigma, C8267) with 10% alcohol. The littermate KO mice and CR mice of the same sex were housed together and received the same handling. Throughout experiments, the researchers were blinded to the genotype. RT-qPCR experiments were used to confirm the efficiency of *Shox2* KO in brains in every animal tested.

In order to view projections of *Shox2*-expressing neurons, we created the Ai27D-*Shox2Cre* mouse. B6.Cg-Gt(ROSA)26Sortm27.1(CAG-COP4*H134R/tdTomato)Hze/J(Ai27D) mice from Jackson labs were crossed with *Shox2*Cre to obtain mice with Shox2-expressing neurons labeled with tdTomato and expressing ChR2.

### Immunohistochemistry

Mice were deeply anesthetized by injection with isoflurane, perfused transcardially with ice-cold PBS followed by 4% paraformaldehyde in PBS and decapitated for brain collection. Mouse brains were placed in 4% paraformaldehyde in PBS at 4◦C overnight for post-fixation. To cryosection the brains, they were sequentially placed in 15 and 30% sucrose in PBS solutions at 4◦C until saturation. The brain samples were embedded in optimal cutting temperature compound and stored at −20◦C and cryo-sectioned in 20–50 μm coronal or sagittal slices with Leica CM3050S cryostat.

For IF staining, slices were washed with 50 mM Tris-Buffered Saline with 0.025% Triton X-100 (TTBS) and blocked in 2% bovine serum albumin (BSA) in TTBS for 2 h at room temperature. Primary antibodies were diluted in blocking solutions and applied on slides overnight at 4^◦^C. Fluorescence-conjugated secondary antibodies were diluted 1:1000 in blocking solutions and applied on slides for 1 h at room temperature. 1:1000 DAPI was applied for 5 min at room temperature for nuclei staining and then washed off. The slices were mounted on slides with mounting media (Vector Laboratories, H-1000) and imaged under a confocal microscope.

### Cytochrome Oxidase Staining for barrels

#### Tissue preparation

The mice were perfused with PBS (0.1M, pH 7.4) followed by 2% PFA. This resulted in softer tissue for flattening which absorbed the CO staining better. Brains were removed and cut down the middle on the sagittal plane with a razor. Using a razor and spatula, subcortical structures were separated from the cortex. Excess cortical regions were removed (i.e., nucleus accumbens, striatum and orbitofrontal cortex) resulting in an even thickness slice of cortex containing the barrels. To ensure smooth flattening, a cut was made across the long side of the frontal cortex. The shelled cortex was placed cortex-down on glass, two rolls of clay (10-20% thinner than brain) were placed on either side to act as spacers, and another glass slide was placed on top. Glass was taped together, and the entire apparatus was placed in a dish of 1% PFA for 24 hours.

#### Slicing and Staining

Cortex was rinsed in PBS (0.1M, pH 7.4) for 15 minutes. Afterwards, 80-150 µM slices were obtained on the vibratome. Slices were rinsed 3 times for 12 minutes in PBS. Staining solution (as described previously) was added, and slices were incubated on a gentle rocker for 3-4 hours at 37℃. The reaction was stopped with one 15-minute rock in 4% PFA. Slices were then washed with PBS 3 times for 10 minutes.

Flattened barrel formation was analyzed using the program Reconstruct. After sectioning, any slice containing a section of barrels was imaged. Images of flattened barrels from the same hemisphere were combined into a project and realigned using the vasculature (visible as white dots/no CO staining within the barrels). Markers for the same vasculature holes were placed on each image, and these were used to align and overlay the images, creating a complete map of flat barrels for each hemisphere.

### RNA Analysis

To confirm knock-down of *Shox2* in the *Gbx2^CreERt/+^* animals, adult *Gbx2^CreERt/+^; Shox2 ^-/f^* male and female mice were anaesthetized by isoflurane inhalation followed decapitation. The brains were removed and a 1 mm thick slice through the thalamus was removed via razor blade, the location of cut is determined by Paxinos and Franklin Mouse Brain Atlas under a stereo-microscope. Medial thalamus tissue is collected with 1mm stainless steel punching tool and lateral thalamus was separated via razor blade. The collected tissues were stored in 50 μL RNA later solution and stored in -80 freezer. To homogenize collected tissues, 350 μL of RLT lysis buffer from Qiagen RNeasy Mini Kit is added to the tissue and homogenized with a pestle mortar. The homogenized tissues went through sonication with a Q55 sonicator (Qsonica) and then 350 μL cold, 70% EtoH was added to the sample. After this, the mixed solution is processed by series of spin and wash follow the instructions book from Qiagen RNeasy Mini Kit. Once RNAs are isolated from tissues, we applied qRT-PCR with Shox2 primer (below) and normalized with GAPDH.

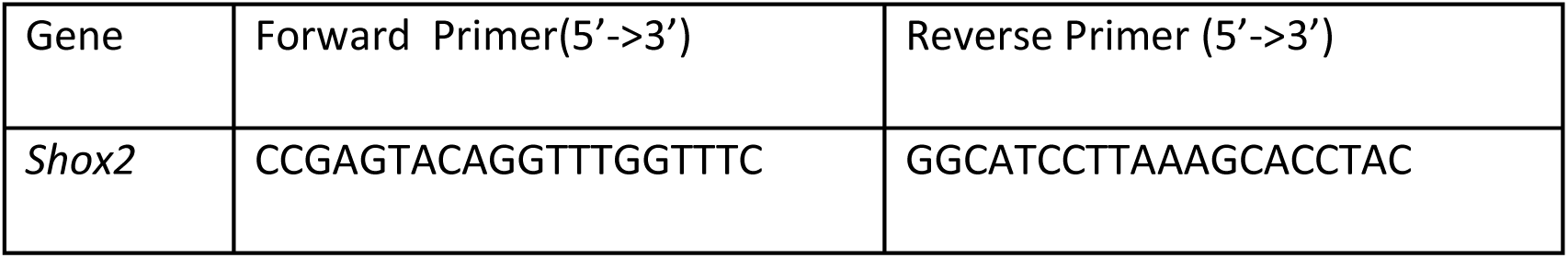

### Electrophysiology

#### Electrophysiology slice preparation

Mice were anaesthetized with isoflurane and rapidly decapitated. Brains were quickly removed and immersed in oxygenated (95% O_2_ and 5% CO_2_), frozen to a slurry, sucrose cutting solution (in mM: 252 sucrose, 2.5 KCl, 26 NaHCO_3_, 1.25 NaH_2_PO_4_·H_2_O, 10 Glucose, 1 CaCl_2_, 5 MgCl_2_). 400µm coronal slices that contained the largest cross-sections of the VB nucleus were obtained with a Vibratome Series 3000 Plus Tissue Sectioning System. The collected slices were transferred to 37°C oxygenated artificial cerebral spinal fluid (aCSF: in mM, 125 NaCl, 2.5 KCl, 26 NaHCO_3_, 1.24 NaH_2_PO_3_, 25 Glucose, 2 MgSO_4_, 2 CaCl_2_) for 20 minutes. Slices were then placed in oxygenated, room temperature aCSF for at least 45 minutes before recordings started. On the electrophysiology rig, slices were continuously perfused with oxygenated, 32°C aCSF. *Shox2*-expressing TCNs were identified by their location within the VB and their expression of RFP, whereas *Shox2*-negative TCNs were identified by location and absence of RFP expression. Glass pipettes were pulled to a resistance of 3-5 MΩ, filled with internal solution (in mM, 120 Kglu, 20 KCl, 0.2 EGTA, 10 Hepes, 4 NaCl, 4 Mg^2+^ATP, 14 phosphocreatine, 0.3 Tris GTP—pH adjusted to 7.2-7.25 by KOH, osmolarity 305-315mOsm) and placed on a silver chloride electrode attached to a headstage. The headstage output to a MultiClamp 700B amplifier, Digidata 1322A digitizer, and a PC running Clampex 10.3 software (Molecular Device).

To determine the effect of *Shox2* expression on the HCN activity in *Shox2*-expressing versus *Shox2*-negative TCNs, the following experiments were performed:

#### Sag

In current clamp, cells were hyperpolarized by -350 pA current with consecutive steps, increasing by +10 pA increments for one second. To calculate sag, which is a gradual depolarization of the membrane while TCNs are hyperpolarized and is generally attributed to the HCN current, the difference between the membrane voltage at the end of the one second hyperpolarization and the beginning of the hyperpolarization step was calculated.

#### HCN current

In voltage clamp, the HCN current was elicited by hyperpolarizing the cell membrane to -150mV for 1 second with incremental steps increasing by 20pA from -50 mV. The amplitude of HCN current will be measured as the difference between the end current of one second hyperpolarization and the beginning instantaneous current at -150mV hyperpolarization.

To determine the effect of *Shox2* expression on the post-anodal burst in *Shox2*-expressing versus *Shox2*-negative TCNs, the following experiments were performed:

#### Area under the curve

In current clamp, a series of one second hyperpolarizing steps, beginning at - 350pA and increasing by 40 pA for 9 traces, were run. The burst following each hyperpolarization was analyzed for the area under the curve. This was done by selecting the section of the trace immediately following the release of hyperpolarization from the moment the membrane potential crosses resting of the neuron until 300 ms after that moment. The entire trace was zeroed to the resting membrane potential of the neuron, and then integrated. The area was then plotted against the stimulus and a two-way ANOVA to determine effects of expression, *Shox2*-expressing versus *Shox2*-negative TCNs, was used to investigate differences.

#### Number of action potentials per post-anodal wave

On the same traces, event analysis was performed with a threshold set to 20mV above the burst wave. Action potentials were considered as events that passed this threshold.

To determine the effect of *Shox2* expression on *membrane properties* in Shox2 TCNs, the following properties were investigated as described:

#### Input resistance

In current clamp, an IV protocol (a series of currents were injected for one second, beginning at -350pA and increasing by 40pA each sweep until it reaches 500pA) was performed. An IV curve (plot of membrane voltage versus current applied) was generated, and input resistance was defined as the slope of the linear portion of the line (approximately the section from -80mV to 40mV membrane potential).

#### Membrane potential

Membrane potential was determined to be the average of a 30 second trace during a time of no membrane activity and while the cell is not being injected with current. Based on recordings thus far, TCN resting membrane potential averages around -70mV, thus, TCNs with a RMP less than -55mV were not considered for analysis.

### Field recordings

Thalamocortical orientation slices were obtained (Agmon & Connors, 1991) from 3-4 week old Shox2Cre;TdtChrd2 mice and maintained in the previously described solutions/recording apparatus. A recording electrode of 1-2 MΩ was filled with aCSF and placed in the layer IV barrel cortex of the slice. A blue LED light was placed as close as possible to the recording electrode, while a 5ms light stimulus was triggered by an external stimulation box.

### Stereotactic surgeries

Prenatal KO of *Shox2* is lethal, thus, a postnatal induction of *Shox2* KO must be utilized. To generate a targeted VB KO mouse model, bilateral VB injections were done to P21 Shox2^fl/fl^ littermates with either a GFPcre virus to generate and label *Shox2* KO cells, or a nonspecific RFP virus to generate and label control cells. After surgery, a one-week incubation period for allowed the virus to take effect and then euthanized the mice at P28 for electrophysiology.

#### Surgeries

Stereotaxic surgeries were performed on P21, Shox2^fl/fl^ mice. Mice were anesthetized with isoflurane, with an oxygen flow rate of 1 L/min and an isoflurane concentration of 2-3% for induction, and 0.5-1.5% for maintenance. Once fully anesthetized, as determined by the lack of response to toe pinch, tail pinch, and eye poke as well as a slowed respiratory rate, ophthalmic ointment was applied to both eyes and subcutaneous buprenorphine was injected into the scruff. Next, mice were ear-barred and secured within a stereotaxic surgery rig. To remove hair from the surgery region, depilation cream was applied, rinsing within 60 seconds. Sterile ethanol wipes and iodine were used to prep the surgery site before a small, lateral incision to the center of the head was made with sterilized scissors and forceps. Forceps and a sterilized razor were used to clear the skull of additional tissue layers. Bregma and lambda were located and aligned to within 0.02 horizontal mm of each other. Then, preloaded pipettes of virus in the nanoject injector were used to find the location of holes to be drilled on left and right sides of the skull. Equal volumes (300nL of approximately 7x10^-12^ titer virus in saline, diluted at a 1:1 ratio) of control and KO virus were injected into P21 VB location (AP: 1.1, ML: 1.5, DV: 3.4; as determined previously by ink injections). Surgery sites were closed using vetbond, and mice are allowed to recover on a heating pad. 1-2 weeks later, mice were used in electrophysiology experiments.

### Ultrasound injections

#### Viral ultrasound injections

P6 mice are anaesthetized via a 4-minute exposure to ice. Beveled, glass pipettes are loaded with virus and attached to a nanoject. Mice are placed on a surgery platform and to stabilize their head, clay is placed on the side opposing the injection. A veterinary ultrasound (VisualSonics) system (with RVM Scanhead model 707B) is used to visualize brain structures and pipette location while the pipette is inserted through the skull and into the VB of the thalamus. 220 nL of virus are injected unilaterally into the VB, and a dwell period of 5 minutes is observed to allow diffusion and diminish capillary expulsion of virus before the pipette is removed from brain. Mice are warmed with a heat lamp to facilitate them regaining consciousness and are then returned to their mother.

### mRNA extraction and qPCR

#### RT-qPCR tissue processing and analysis for mRNA expression

To confirm knockdown of *Shox2* in the *Shox2*^fl/fl^+ viral AAV2cre model, unilateral injections were accomplished. The GFPcre virus was injected into the left VB of Shox2^fl/fl^ mice, and control, non-specific RFP virus into right VB, creating a within mouse control with *Shox2* KO on the left and control conditions on the right. Mice were assessed at P34 for *Shox2* mRNA expression. Unilateral KO male and female mice were anesthetized by isoflurane inhalation followed by decapitation. The brains were removed and a 1-mm thick slice through the thalamus was removed via razor blade, the location of cut was determined by Paxinos and Franklin Mouse Brain Atlas. VB thalamus tissue was collected with 1-mm stainless steel punching tool. The collected tissues were stored in 50 μL RNA later solution and stored in −80 freezer. To homogenize collected tissues, 350 μL of RLT lysis buffer from Qiagen RNeasy Mini Kit was added to the tissue and homogenized with a pestle mortar. The homogenized tissues went through sonication with a Q55 sonicator (Qsonica) and then 350 μL cold, 70% EtoH was added to the sample. RNA was extracted according the instructions (Qiagen RNeasy Mini Kit), and qRT-PCR was performed with the *Shox2* primer described above and normalized with GAPDH (F: GTCGGTGTGAACGGATTTG, R: TAGACTCCACGACATACTCAGCA) to determine expression of *Shox2* mRNA.

### EEG implantation and spindle frequency detection

Animal experiments were conducted in accordance with approved IACUC guidelines at Baylor College of Medicine. To record EEG activity, WT and *Shox2* KO mice (age P90) were implanted with subdural stainless-steel screw and intracranial electrodes (PlasticsOne) using aseptic surgical technique. Two hours prior to surgery, mice were given sustained-release buprenorphine (1mg/kg b.w.). Mice were anesthetized with isoflurane (5% induction, 1-2% maintenance) and fixed to a stereotaxic frame with a heating pad for thermal support. Hair was removed from the surgical site with clippers and depilation cream and then cleaned with three alternating swabs of betadine scrub and alcohol. A local injection of lidocaine/bupivacaine block was given and a 1-2cm midline sagittal cut was made along the scalp. After removal of periosteum from the skull, three small burr holes were drilled at the following coordinates (Medial-Lateral, Anterior-Posterior, Dorsal-Ventral): somatosensory cortex (3mm, -1.5mm, 0.5mm), motor cortex (1.5mm, 1.5mm, 0.5mm), and reference electrode (-1.5mm, 1.5mm, 0.5mm). A smaller hole was drilled for an intracranial straight wire electrode to record from VB thalamus (1.5mm, -1.5mm, 3.5mm). A spring wire electrode was sutured into the paraspinous cervical muscle to record electromyogram (EMG) activity. The screw and wire electrodes were coupled to pins and were connected to a headmount secured with Metabond and dental cement. Mice were singly housed and allowed to recover for 3 days following surgery. After recovery, mice were connected via tether to a commuter for continuous video-synchronized EEG (vEEG) recording in a cylindrical chamber with food and water (35.5cm D x 30.5cm H; Pinnacle Technology Inc.). vEEG activity was acquired with a Nicolet v32 amplifier (2 kHz sampling) and an overhead digital camera synched to Natus EEG acquisition software v9.54 (Natus, Pleasanton, PA). Post-acquisition artifact removal and processing was conducted offline with Labchart V8 software (AD Instruments, Colorado Springs, CO). vEEG activity was analyzed blind to genotype. To determine spectral power, raw EEG signals were bandpass filtered at 0.5-50Hz in LabChart and converted to the frequency domain using fast Fourier transform with the Hann (cosine-bell) method and 2000 FFT size with 50% window overlap (Martinez et al, 2020; Born et al, 2021). EMG was bandpass filtered at 50-200Hz to optimize classification of vigilance states. Five-minute epochs, free of noise and muscle artifact, were selected for spectral power analysis. Sleep scoring was carried out for a full 24hr light/dark cycle (14hr/10hr) by examining EEG traces and EEG spectral heat maps. Wake/sleep transition states were confirmed by visual inspection of video recordings. Awake states were defined as high amplitude EMG/low amplitude, high frequency EEG activity. Sleep states were segmented into slow-wave sleep (non-REM) and REM sleep. Non-REM was defined by low amplitude EMG/high amplitude, low frequency EEG activity. REM sleep was defined by low amplitude EMG/low amplitude, high frequency EEG activity. Sleep duration and spectral power were quantified in each sleep state during light and dark cycles. Sleep spindles were detected by filtering the EEG signal at the sigma frequency as previously described (Copping & Silverman, 2021; Kim et al., 2015) Briefly, spindles were detected from S1 and VB EEG signals in Labchart by first applying a bandwidth filter in the sigma frequency (9-16Hz), then applying a Butterworth filter, followed by obtaining the root mean square in 750ms square windows. The resulting filtered signal was then cubed to reduce background noise and amplify sigma frequency peaks. Spindles were quantified using the Peak analysis function in Labchart. Due to variability in spindle amplitude, spindle threshold was determined empirically for each mouse, however, mean threshold between genotypes did not differ (WT 97.5mV ± 44.7µV and KO 52.3mV±30.4µV, p>0.05) Total spindle frequency was quantified during the light cycle. Spindle frequency was also compared during the one minute immediately before the state transition from non-REM to REM sleep and separately during the transition from non-REM to the wake state.

### Behavior

#### Adhesive tape task

Mice were habituated to the recording chamber for 3 hours, 3 consecutive days in a row. On the third day, after 2 hours, videos were recorded as stickers were placed on either left-or right-side whiskers. Latency to removal was scored by a blinded observer. Each value is an average of three trials.

The paw sensation test was applied to assess mouse paw sensorimotor response (Arakawa et al., 2014). A small piece of round sticky paper tape (Tough-spots, for microtube cap ID, ∼1cm2, Research Products International 45 Corp. 247129Y) was applied to the plantar surface of the right hindpaw of each mouse and the mouse was placed back to its home cage. The latency to the first response to the paper of each mouse was measured and analyzed

### Open field test

Open field test was applied to test mouse exploratory behavior and anxiety behavior. All mice have received routine handling for a week and have put in the same room for 1 hour each day for three days before experiments. Each mouse was place in the open field and allowed to explore freely under dim red light for 5 minutes. Infrared beams and computer-based software Fusion was used to track mice and calculate mice activity and time spent in the center (4x4) of total open field (8x8).

### Novel object recognition

In novel object recognition (NOR) experiments, all mice have received routine handling and three days room habituations before experiments. On the 44 day before familiarization trial, each mouse was placed in the open field in the absence of objects and allowed to explore it freely for open field habituation, the behaviors of which were recorded and analyzed further as open field test data. In the familiarization trial, each mouse was placed in the open field containing two identical 100 ml beakers in the neighboring corners for 5 minutes. Twenty-four hours later, each mouse was placed back in the same open field with two objects, one of which was the identical 100 ml beaker and the other one was replaced with a novel object (a lock in a similar dimension) for 5 minutes testing trial. To prevent coercion to explore the objects, mice were always released against the center of the opposite walls in both familiarization and testing trials. The mouse behaviors in the testing trials were taped and analyzed by experimenters blinded to the genotypes of the mice. The exploration was defined as the nose of the mouse is sniffing and touching it with attention, while running around the object, sitting or climbing on it was not recorded as exploration (Antunes and Biala 2012).

